# Comparative studies of transmission mode and localisation patterns of common RNA viruses in Queensland fruit fly (*Bactrocera tryoni*) reveal most are vertically transmitted

**DOI:** 10.64898/2026.03.20.713308

**Authors:** Farzad Bidari, Jennifer L. Morrow, Sanjay Kumar Pradhan, Markus Riegler

## Abstract

RNA viruses are common in tephritid fruit flies including the Queensland fruit fly, Australia’s most significant horticultural pest. For many their transmission, tissue tropism and load across host development remain unexplored. Yet these factors are important for host biology, ecology and pest management. We investigated Bactrocera tryoni orbivirus (OV), Bactrocera tryoni xinmovirus (XV), Bactrocera tryoni toti-like virus (TLV) and Bactrocera tryoni iflavirus species 2 (IVsp.2) that commonly coinfect *B. tryoni* laboratory populations. OV and XV transmission was vertical within and on eggs, while TLV transmission was vertical within eggs. IVsp.2 was not detected in eggs but was present in adults; however, IVsp.2 was horizontally transmitted, with viral load increasing with cohabitation time with infected flies. Horizontal transmission was not observed for the other viruses. OV had a similar load across all tissues, while XV was consistently more abundant in ovaries. TLV had a high viral load in the brain whereas IVsp.2 was abundant in the thorax, foregut and midgut. Besides differences in eggs, the viruses were detected in all other developmental stages, but viral load patterns differed: viral load remained constant for TLV, fluctuated for OV and XV, and was low in pre-adult stages and high in adults for IVsp.2. Our findings demonstrate distinct transmission strategies and tissue tropism among the viruses, providing new insights into their epidemiology and role in host biology. Furthermore, contrary to prevailing views that viruses are generally horizontally transmitted, most known RNA viruses of *B. tryoni* are vertically transmitted affecting the evolution of host-virus interactions.

## 1. Introduction

RNA viruses are diverse, contain RNA as their genetic material and include many of the most rapidly evolving and ecologically significant viruses (Koonin et al., 2006; Shi et al., 2016). Transcriptome studies of insects have detected a large number of RNA viruses within them, revealing that insects can serve as hosts and reservoirs of diverse RNA viruses (Shi et al., 2016; Li et al., 2015). Numerous RNA viruses have been detected in the transcriptomes of true fruit fly species (Tephritidae) which include about 350 pests of economic importance (PHA, 2018; White et al., 1992). One transcriptome study of nine species of the tephritid genera *Ceratitis*, *Bactrocera* and *Zeugodacus* detected over eight RNA virus families, with some viruses occurring in several tephritid species (Sharpe et al., 2021). Other studies focusing on individual tephritid fruit fly species revealed 13 RNA virus species belonging to nine families in the Mediterranean fruit fly *Ceratitis capitata* (Hernández-Pelegrín et al., 2022), 11 RNA viruses species in the oriental fruit fly *Bactrocera dorsalis* (Zhang et al., 2020; Zhang et al., 2022) and ten RNA virus species for the melon fly *Zeugodacus cucurbitae* (Pradhan et al., 2024a).

Queensland fruit fly, *Bactrocera tryoni*, is Australia’s most important horticultural pest. It originates from the rainforests of tropical and subtropical coastal Queensland and northern New South Wales and is now widely distributed across northern and eastern Australia, as well as in parts of the Pacific region where it can infest a wide range of fruits and fruiting vegetables (Dominiak and Daniels, 2012; Hancock et al., 2000; Popa-Báez., 2021). It hosts several RNA viruses including Bactrocera tryoni cripavirus (CV; *Dicistroviridae*), Bactrocera tryoni iflavirus sp. 1 and Bactrocera tryoni iflavirus sp. 2 (IVsp.1 and IVsp.2; *Iflaviridae*), Bactrocera tryoni sigmavirus with the ICTV-recognised name *Sigmavirus tryoni* (SV; *Rhabdoviridae*), Bactrocera tryoni toti-like virus (TLV; *Betatotivirineae*), Bactrocera tryoni orbivirus (OV; *Sedoreoviridae*) and Bactrocera tryoni xinmovirus with the ICTV-recognised name *Trocevirus haikouense* (XV; *Xinmoviridae*), all detected in transcriptomes of laboratory flies (Sharpe et al., 2021; Sharpe et al., 2026). Five of these viruses (CV, IVsp.1, OV, TLV and XV) were found to coinfect *B. tryoni* laboratory populations across multiple generations without showing obvious symptoms, suggesting that they can establish persistent covert infections (Sharpe et al., 2021; Sharpe et al., 2026). However, a comparison of a fly population coinfected with these five viruses had slower egg-to-pupa development, lower emergence and lower adult survival under stress than a genetically related fly population established with flies of the same source population lacking CV and IVsp.1 but coinfected with the other three viruses (Sharpe et al., 2026). Furthermore, the injection of flies with a mixed suspension containing these five viruses significantly increased mortality, with IVsp.1 being the only virus that increased in viral load suggesting that it is responsible for the increased mortality (Sharpe et al., 2026).

Similar persistent covert infections with RNA viruses have been reported in other tephritids and insects, where their effects on host fitness often depend on environmental or physiological conditions (Llopis-Giménez et al., 2017; Virto et al., 2017). Studies in the vinegar fly (*Drosophila melanogaster*), the western honey bee (*Apis mellifera*) and the beet armyworm (*Spodoptera exigua*) further show that covert virus infections can become pathogenic under stress or subtly influence immunity, reproduction and behaviour, leading to ecologically significant outcomes (Gupta et al., 2017; Remnant, 2019; Mengual-Marti, 2022).

In general, RNA viruses can be transmitted vertically, horizontally or via a combination of both vertical and horizontal transmission (Hajek and Shapiro-Ilan, 2018; Solter and Becnel, 2017; Cory, 2015). Vertical transmission can be maternal either within the egg (transovarial or transovarian) or on the surface of the egg (transovum), or paternal through sperm or seminal fluid, within the egg or on the surface of the egg. Horizontal transmission can occur via food, faeces or wounds (Hajek and Shapiro-Ilan, 2018; Solter and Becnel, 2017; Cory, 2015). Recent studies on virus transmission in *B. tryoni* have found that CV is transmitted horizontally, IVsp.1 predominantly maternally (within the egg), and SV biparentally (also within the egg) albeit more effectively maternally than paternally (Morrow et al., 2023; Pradhan et al., 2026). Previously, viruses were considered to spread predominantly through horizontal transmission, including direct contact, vector-mediated routes or environmental exposure (Fermin, 2018). However, an increasing number of studies in tephritid fruit flies, including our previous studies on *B. tryoni* (Morrow et al., 2023; Sharpe et al., 2024; Pradhan et al., 2026) and recent studies on *C. capitata* (Hernández-Pelegrín et al., 2024b; Longdon et al., 2017) indicate that vertical transmission is the predominant mode of transmission for most RNA viruses for which transmission modes have been investigated in tephritids. However, the transmission modes of many other RNA viruses of *B. tryoni* and their localisation across tissues and developmental stages remain unclear.

This knowledge gap applies to many if not most viruses detected in insect and other arthropod transcriptomes. Filling this gap is critical, as these parameters influence the persistence of RNA viruses and their effects on hosts, including in arthropods that are mass-reared for various purposes, such as feed, food and silk production (e.g. black soldier fly, silkworm), pollination services (e.g. honey bee), as well as biological control (e.g. parasitoid wasps and predatory mites) (Maciel-Vergara and Ros, 2017; Bertola and Mutinelli, 2021). It is similarly important to investigate the biology and epidemiology of these viruses for improving and ensuring the quality of insects used in sterile insect technique (SIT) programs (Hendrichs and Robinson, 2021; Fisher, 2020).

In general, vertically and horizontally transmitted viruses may have different developmental stage and tissue tropism. Specifically, vertically transmitted viruses may be present across all developmental stages, including eggs and embryos as the key link for vertical virus transmission between generations. In contrast, horizontally transmitted viruses may be acquired by the physiologically more active larval and adult stages in which they may replicate whereas horizontally transmitted viruses should be absent from eggs and embryos. Therefore, vertically transmitted viruses may be detected across all tissues particularly in reproductive tissues, whereas horizontally transmitted viruses may target metabolically active tissues such the digestive tract and muscle tissues (Morrow et al., 2023; Ponnuvel et al., 2022; Carrillo-Tripp et al., 2014; Hernández-Pelegrín et al., 2024a; Kang et al., 2022).

Here we investigated the transmission modes of the four viruses OV, XV, TLV and IVsp.2 using *B. tryoni* laboratory populations that either carry these viruses as persistent covert infections or are free of these viruses. Additionally, the viral load of these viruses was assessed across different developmental stages and in various somatic and reproductive tissues in order to investigate infection patterns for vertically versus horizontally transmitted viruses.

## 2. Materials and methods

### 2.1 *Bactrocera tryoni* laboratory populations

We used five laboratory populations of *B. tryoni* (C28, GOS, HAC, LE, UNSW) and one laboratory population of *Bactrocera jarvisi* (Bj) (Table S1). The populations were reared as previously described (Morrow et al., 2015) and maintained in separate Bugdorm cages (30 cm × 30 cm × 30 cm) at 25 °C ± 3 °C, 65% ± 15% relative humidity and natural light. To prevent virus cross-contamination, all cages, food and water containers were sterilized with standard household bleach (4%) and thoroughly rinsed before use.

### 2.2 RNA extraction and cDNA synthesis

RNA was extracted from individuals for the assessment of virus prevalence and load in different developmental stages, and from pooled samples for the assessment of virus load in eggs and different tissues. RNA extraction was performed using 500 µl TRI Reagent (Sigma-Aldrich) according to the manufacturer’s instructions, with the RNA resuspended in 30 μL of nuclease-free water. The RNA was then used for cDNA synthesis using the RevertAid First Strand cDNA Synthesis Kit (Thermo Scientific), employing equal amounts of random hexamers and oligo d(T)-18 primers (Thermo Scientific).

### 2.3 RT-PCR and RT-qPCR

Virus detection was carried out using reverse transcription PCR (RT-PCR) and RT quantitative PCR (RT-qPCR) with cDNA as the template. Virus-specific primer sets targeting the viral RNA-dependent polymerase (RdRp) genes of OV, XV, TLV, IVsp.2, IVsp.1, CV and the nucleocapsid (N) gene of SV were used for virus detection and quantification (Table S2) (Sharpe et al., 2021; Sharpe et al., 2026). The efficiency of all primer sets ranged between 91% and 107% as determined by amplifying tenfold serial dilutions of cDNA samples harbouring high loads of each virus (Supplementary File). This was also used to determine that the amplification threshold for each primer set was 33 to 34 cycles. For SV a N gene primer set was used because sigmavirus genes located closer to the 3′ end, particularly the N gene, may be present at higher transcript abundance (Dietzgen et al., 2017). Also, to minimize pipetting errors and ensure optimal template concentration, the cDNA was diluted with water (1:4.2) prior to PCR.

RT-PCR reactions were performed in 10 μL containing 5x MyTaq Red Reaction buffer (Meridian Bioscience), 0.25 U MyTaq DNA polymerase (Meridian Bioscience), 5 μM of each primer and 2 μL of diluted cDNA. Thermocycling conditions were 94 °C for 3 minutes, followed by 34 cycles of 94 °C for 1 minute, 54 °C (OV, XV, TLV, IVsp.2 and IVsp.1) or 58°C (CV) for 30 s and 72°C for 1 minute, with a final 10 minutes at 72 °C. The sets of diagnostic RT-PCRs were conducted on individual flies to evaluate the virus prevalence in laboratory populations. Ten males and females (n=20) were randomly collected from each of the six laboratory populations and RT-PCR was performed on their cDNA as described above. To confirm primer specificity and virus identity, RT-PCR amplicons were Sanger sequenced using the BigDye Terminator v3.1 Cycle Sequencing Kit (Thermo Fisher Scientific). Before this, the amplicons were ExoSAP-treated with 0.25 U exonuclease I (New England Biolabs) and 0.25 U TSAP thermosensitive alkaline phosphatase (Promega) at 37 °C for 30 minutes, followed by 95°C for 5 minutes.

RT-qPCR reactions were performed in 10 μL containing 5 μL SensiFAST SYBR (Millenium Sciences), 0.4 μL of each 10 μM primer and 4.2 μL diluted cDNA template. Thermocycling conditions were 95 °C for 3 minutes, followed by 35 cycles of 95 °C for 5 s, 58 °C for 10 s and 72°C for 20 s. The PCR run ended with a disassociation curve step in which the temperature increased from 50°C to 98 °C in 1 °C increments every 5 s.

Each gene was tested in duplicate and dissociation curves were reviewed for each reaction. A sample was deemed virus-positive if the melt-curve analysis of the amplicon showed a single peak at the correct temperature (±0.5 °C) with minimal primer dimer formation and a Cq value of 35 or less, where Cq values of technical replicates differed by at most 1.5 cycles. Samples that did not meet these criteria for both technical replicates were classified as non-amplification. Furthermore, any technical variation was controlled for by the biological replication of three to five, depending on the type of experiment.

Diagnostic RT-qPCRs were performed with normalisation against the single-copy host reference gene elongation factor 1 alpha (*ef1a*) which had previously been found to be stably expressed across different host stages (Morrow et al., 2023). Viral Cq values were normalised to *ef1a* Cq values using the formula 2^-(ΔCq)^ (samples without amplification were assigned a Cq value of 35, equivalent to the maximum number of PCR cycles and below the detection threshold of 33-34 cycles, to allow normalisation of these samples and log transformation for visualisation). As all primer efficiency values were between 91% and 107%, no adjustment for efficiency was required (Schmittgen and Livak, 2008). The minimum threshold for categorizing virus infection was established at one viral transcript per ten million *ef1a* transcripts (10e-7).

### 2.4 Testing and surface bleaching of eggs to assess vertical transmission

To test for vertical transmission of viruses via eggs, 30 mated females (approximately four weeks old) each of HAC, GOS and C28 were placed in three separate cages with sugar, water and yeast hydrolysate mix (Morrow et al., 2023) for one week. An oviposition cup containing orange juice and covered with punctured parafilm was provided for three hours. Then, 360 unhatched eggs were collected from each cage. Eggs were divided into four treatment groups, each with three replicates (30 eggs per replicate). The control group remained untreated, while the other three groups were placed in 4% sodium hypochlorite (Sigma-Aldrich) for 30 s, 2 minutes or 5 minutes, followed by thorough rinsing in fresh DNase and RNase-free distilled water (Morrow et al 2023; Pradhan et al 2026). Surface bleaching of eggs may remove viruses that may be present on the outer membrane (chorion) and be transmitted outside the eggs. In contrast, viruses transmitted within eggs will remain detectable after surface bleaching (Morrow et al 2023; Pradhan et al 2026). For each cage RNA was extracted from each of the 12 pooled samples (30 eggs each) and from eight randomly collected female flies. RT-qPCR for viral load assessment was then performed as described above.

### 2.5 Cohabitation of females of infected and uninfected fly populations to evaluate horizontal transmission

For the cohabitation test to assess horizontal virus transmission, we used pairs of infected and uninfected fly populations. For the XV and OV horizontal transmission experiments we used C28 (infected with XV and OV) and LE (free of XV and OV); for the IVsp.2 horizontal transmission experiments LE (infected with IVsp.2) and UNSW (free of IVsp.2); and for the TLV horizontal transmission experiments C28 (infected with TLV) and Bj (free of TLV), noting that, because all *B. tryoni* laboratory populations had TLV, two fly species were used for the TLV horizontal transmission experiment, i.e., *B. tryoni* and *B. jarvisi*. Albeit the two are distinctly different species they are closely enough related so that they can hybridise under laboratory conditions (Morrow et al., 2014). For each population pair, three cages were set up, each containing 15 virgin females from both the infected and uninfected populations, and subjected to three different cohabitation periods: one day (cage 1), five days (cage 2) and ten days (cage 3). After each of these cohabitation periods, five females of the exposed uninfected populations and nine females of the infected populations were individually placed into microcentrifuge tubes and stored at −80 °C. Females were identified by the difference in eye colour between LE (a mutant line with lemon-coloured eyes) and the other wild-type eye-coloured populations (Zhao et al., 2003) and by the morphological differences between C28 (*B. tryoni*) and Bj (*B. jarvisi*) (PHA, 2018). Then, the remaining ten females of the uninfected populations were kept individually in small containers for an additional five or ten days to evaluate virus load after this extended incubation period. Females were then individually placed into microcentrifuge tubes and stored at −80 °C. RNA extraction from individual flies, cDNA synthesis and RT-qPCR were carried out as described above.

### 2.6 Detection of viral load in various tissues

To measure the viral load in different tissues, 15 virgin males and 30 virgin females each of HAC and GOS were collected. Emerged males were kept in one cage and fed sugar and water for one week, while emerged females were separated into two cages (15 flies per cage), one with sugar, water and a yeast hydrolysate mix required for ovary development (Morrow et al., 2015) and the other with only sugar and water for one week. After this, the flies were individually dissected in fresh DNase and RNase-free distilled water under a stereomicroscope to collect the following tissues: brain tissue (after removing the mouthparts, antennae and head exoskeleton), salivary glands, thorax, foregut plus anterior midgut and ovaries (for females) or testes (for males). Individual tissues were briefly rinsed two times in fresh DNase and RNase-free distilled water prior to pooling to minimise residual virus and potential cross contamination. For each line and adult type, the tissue types of five individuals were immediately pooled in microcentrifuge tubes kept on ice. For viral load analysis by RT-qPCR, RNA was extracted and cDNA synthesised for three replicates (each with the pooled tissues of five individuals) for each tissue and adult type.

### 2.7 Detection of viral load across developmental stages

Viral load was assessed across five developmental stages (eggs, larvae, pupae, adult males and females) for HAC and GOS. Four-week-old (post-eclosion) females were allowed to oviposit into small cups containing larval diet covered with punctured parafilm (Morrow et al., 2015). For each of the two populations, five fully developed third instar larvae that had exited the larval diet for pupation after 8-10 days, five pupae (three days after pupation) and five virgin males and five virgin females (less than one day after emergence) were individually placed into microcentrifuge tubes and stored at −80 °C. Furthermore, a total of 120 eggs from each population were collected from another oviposition cup after HAC and GOS flies were allowed to oviposit for three hours, separated into four pools of 30 eggs per microcentrifuge tube and stored at −80 °C. RNA was then extracted from individual larvae, pupae, adult males and females, and the pooled egg samples, followed by cDNA synthesis. Viral load was measured using RT-qPCR.

### 2.8 Statistical analyses

Data normality was checked using Shapiro-Wilk and Kolmogorov-Smirnov tests. Fisher’s exact test and Cochran’s test were performed using IBM SPSS (version 30.0.0.0) to analyse the virus prevalence in the laboratory populations. The viral load data across the developmental stages, tissues and transmission experiments were analysed using IBM SPSS (version 30.0.0.0) and R software (2024.12.0+467). One-way ANOVA and Kruskal-Wallis tests were used to test for significance, and if results were significant post-hoc pairwise comparisons were performed using Tukey’s HSD and Dunn’s tests with Bonferroni correction. The significance threshold was set at *p* < 0.05. Graphs were created using GraphPad Prism v8.4.0 and R v4.4.2 (R Core Team, 2024). Plots for virus prevalence and coinfection were constructed using the Mondrian package v1.1.2 (Siberchicot et al., 2025) in R.

## 3. Results

### 3.1 Virus prevalence in different laboratory populations

Coinfections with three to seven of the seven viruses were found in females and males of all six *B. tryoni* laboratory populations (Figure 1). There was no significant difference in virus prevalence between females and males within each population (Fisher’s exact tests, all *p* values > 0.05; Supplementary File), so their prevalence was analysed across both sexes combined. The prevalence of the seven viruses was significantly different across populations (*p* < 0.01) (Table S3). The four focal viruses of our study (OV, XV, TLV and IVsp.2) were observed in both females and males of GOS, HAC and C28 (Figure 1), although OV load in HAC was low (Figure 2a). High coinfection prevalence of TLV and IVsp.2 (without the other two viruses) was observed in LE, while Bj had mostly just IVsp.2, and UNSW had TLV without any of the other three viruses (Figure 1). Besides these four viruses, IVsp.1 was detected exclusively in C28 at a high prevalence; CV was present in all populations at a high prevalence in GOS, LE and C28, moderate prevalence in HAC and UNSW and low prevalence in Bj; and SV was detected in HAC and UNSW at high prevalence, in GOS and LE at moderate prevalence, and in Bj and C28 at low prevalence.

**Figure 1.**
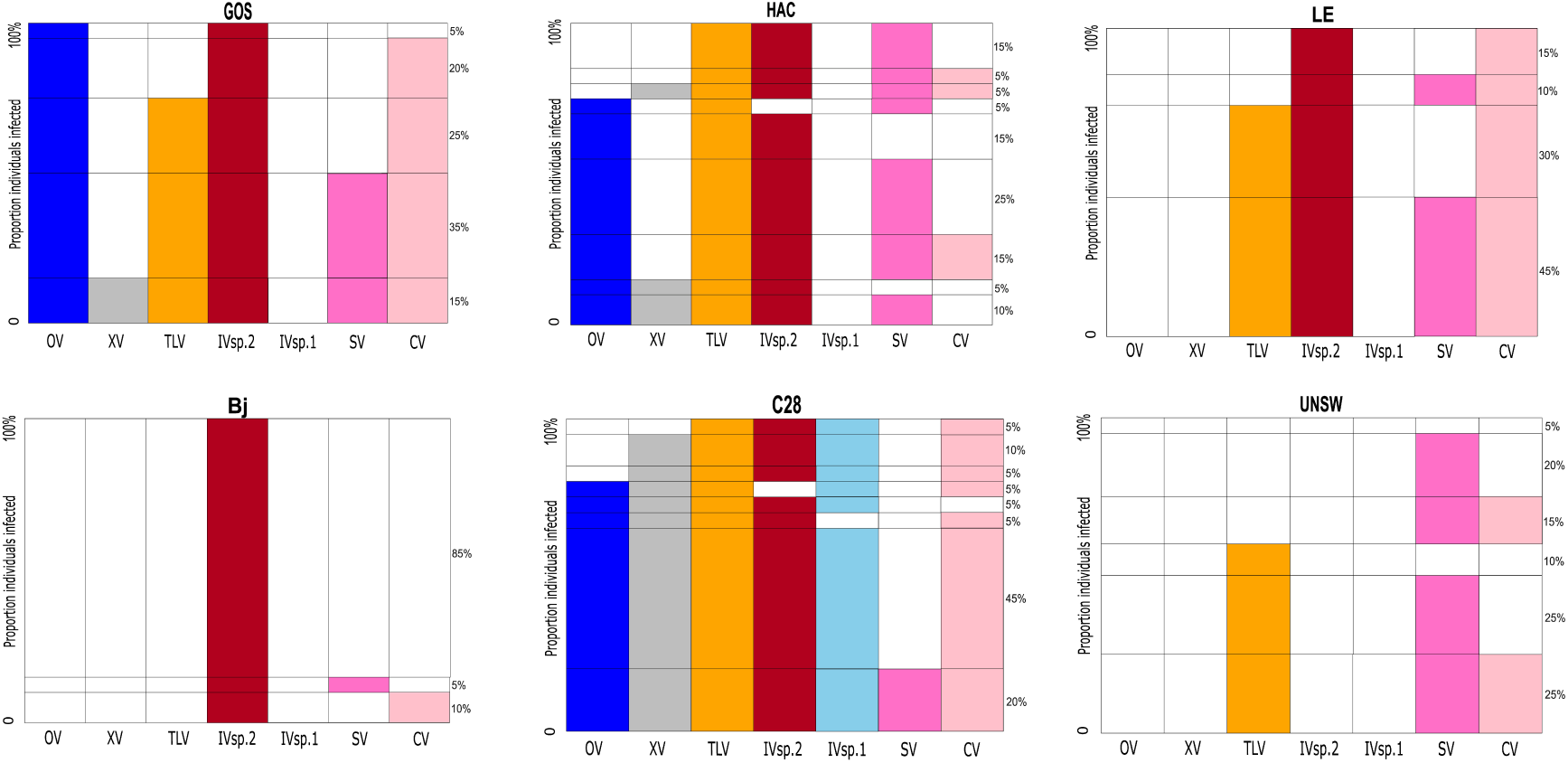
Prevalence of Bactrocera tryoni orbivirus (OV), Bactrocera tryoni xinmovirus or *Trocevirus haikouense* (XV), Bactrocera tryoni toti-like virus (TLV), Bactrocera tryoni iflavirus sp. 2 (IVsp.2), Bactrocera tryoni iflavirus sp. 1 (IVsp.1), Bactrocera tryoni sigmavirus or *Sigmavirus tryoni* (SV) and Bactrocera tryoni cripavirus (CV) in males and females of *Bactrocera tryoni* and *Bactrocera jarvisi* laboratory populations. The plots show for each population the proportion of individuals infected with OV (dark blue), XV (grey), TLV (orange), IVsp.2 (red), IVsp.1 (light blue), SV (pink) and CV (salmon), in (a) GOS, (b) HAC, (c) LE, (d) Bj, (e) C28 and (f) UNSW. Plots are presented as Mondrian (heatmap) plots, where the height of each block within a column represents the percentage of individuals infected with a particular virus combination (indicated on the right side of the plots). Male (n=10) and female (n=10) data were combined as no significant differences in virus prevalence were detected between the two sexes. Viruses are listed with the four focal viruses of this study (OV, XV, TLV, IVsp.2) on the left and previously studied viruses (IVsp.1, SV, CV) to the right whereby within both of these two groups the five vertically transmitted viruses (OV, XV, TLV; IVsp.1, SV) are listed first followed by the two horizontally transmitted viruses (IVsp.2 and CV).

**Figure 2.**
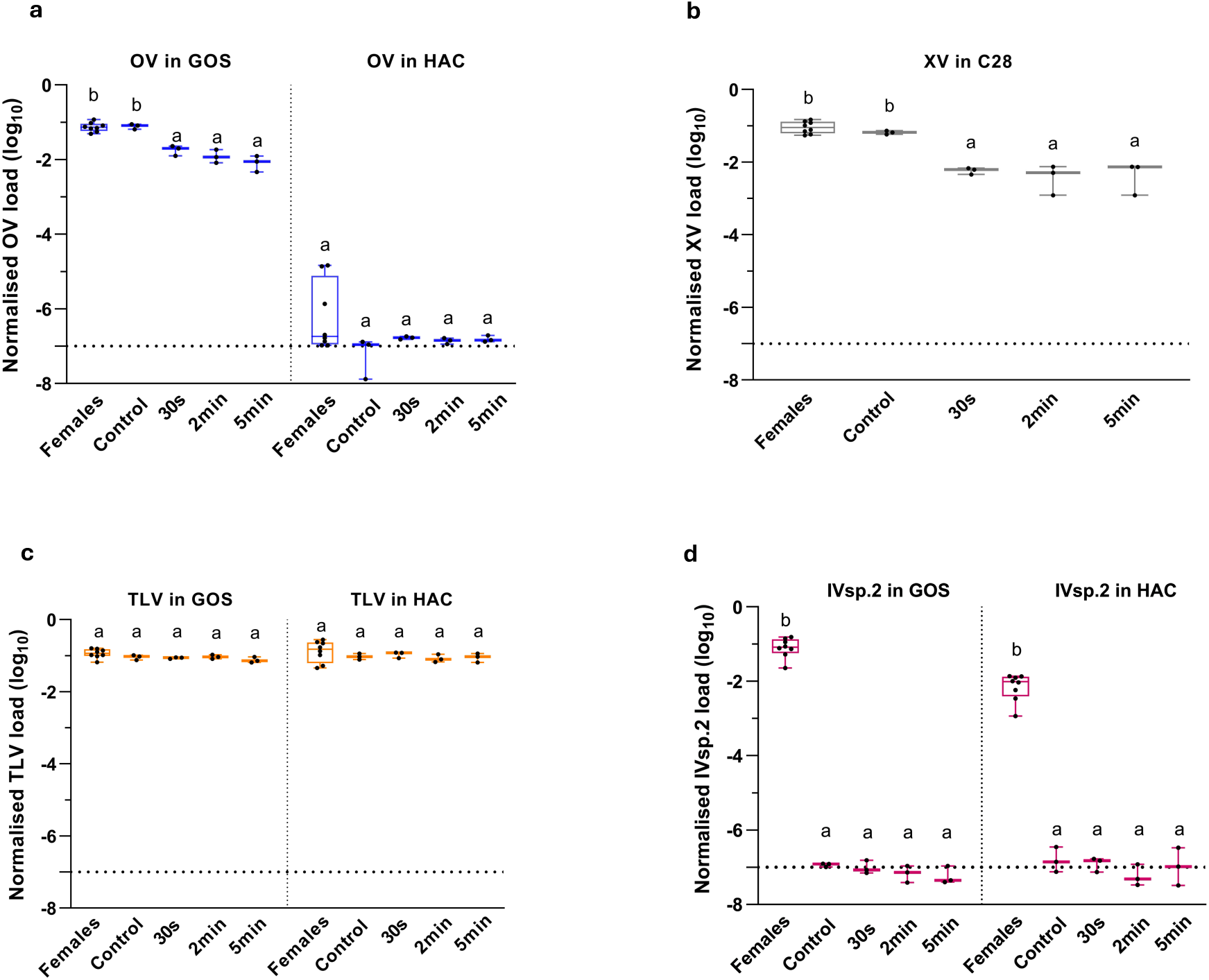
Normalised viral load (log_10_) of (a) Bactrocera tryoni orbivirus (OV), (b) Bactrocera tryoni xinmovirus or *Trocevirus haikouense* (XV), (c) Bactrocera tryoni toti-like virus (TLV) and (d) Bactrocera tryoni iflavirus sp. 2 (IVsp.2) in eggs (control, bleached for 30 s, 2 minutes and 5 minutes) of infected females of GOS, HAC and C28. The virus detection threshold was set at 1e-7 virus-specific RNA-dependent polymerase (RdRp) or nucleocapsid (N) per *ef1a* transcript, shown by the horizontal dashed line. Means that differ significantly from each other are represented by different letters for each population separately.

### 3.2 Egg experiments to assess vertical transmission

OV (Figure 2a) in GOS, and XV (Figure 2b) and TLV (Figure 2c) in both GOS and HAC were detected in females and eggs (control and bleached for 30 s, 2 minutes and 5 minutes). The TLV load was similarly high in both bleached and unbleached eggs, and did not differ significantly between females and eggs in GOS (F₄,₂₀ = 2.476, *p* = 0.089) and HAC (F₄,₂₀ = 1.223, *p* = 0.342) (Figure 2c). In contrast, surface bleaching of eggs significantly reduced the viral load of OV in GOS compared to the control (F₄,₂₀ = 20.07, *p* < 0.01), with a 4.3 fold reduction after 30 s bleaching, when OV still presented a mean normalized viral load of 0.0179; furthermore, the OV load did not differ across the 30 s, 2 and 5 minute bleaching treatments (Figure 2a). For XV in C28, viral load in eggs also decreased significantly after bleaching relative to the control (F₄,₂₀ = 14.642, *p* < 0.01), with an 11.3 fold reduction after 30 s bleaching when it still had a mean normalized viral load of 0.0057; similar to OV in GOS, it remained high and did not differ across 30 s, 2 and 5 minutes (Figure 2b). Unlike OV in GOS, no significant differences were found for OV in HAC across all bleaching treatments. OV in HAC eggs and most females was just above the detection limit; however, in some females, the viral load was higher than in eggs (F₄,₂₀ = 0.865, *p* = 0.5) (Figure 2a). In contrast to the other three viruses, IVsp.2 was only detected in females of GOS and HAC but not in eggs. In both populations, IVsp.2 was at the detection limit in eggs and significant differences in viral load were observed between females and eggs in GOS (F₄,₂₀ = 10.617, *p* < 0.01) and HAC (F₄,₂₀ = 8.520, *p* < 0.01) (Figure 2d). Therefore, OV, TLV and XV showed signs of vertical transmission with OV and XV being found within and on the surface of eggs, while TLV was only within eggs. In contrast, IVsp.2 was not detected in eggs suggesting that it may rely on horizontal transmission.

### 3.3 Cohabitation experiments to evaluate horizontal transmission

Despite varying cohabitation times (1, 5 and 10 days) and incubation times (0, 5 and 10 days), no horizontal acquisition of OV, XV and TLV was detected in females of the uninfected populations exposed to females of infected populations. For OV, a significant viral load difference was observed between females of infected (C28) and uninfected populations (LE) used in the experiment (Kruskal-Wallis chi-squared 34.58, df 11, *p* <0.01) (Figure 3a). A similar pattern was observed for XV in LE flies exposed to infected C28 flies (Kruskal-Wallis chi-squared 56.13, df 11, *p*<0.01) (Figure 3b) and TLV in Bj flies exposed to infected C28 flies (Kruskal-Wallis chi-squared 54.58, df 11, *p* <0.01) (Figure 3c).

**Figure 3.**
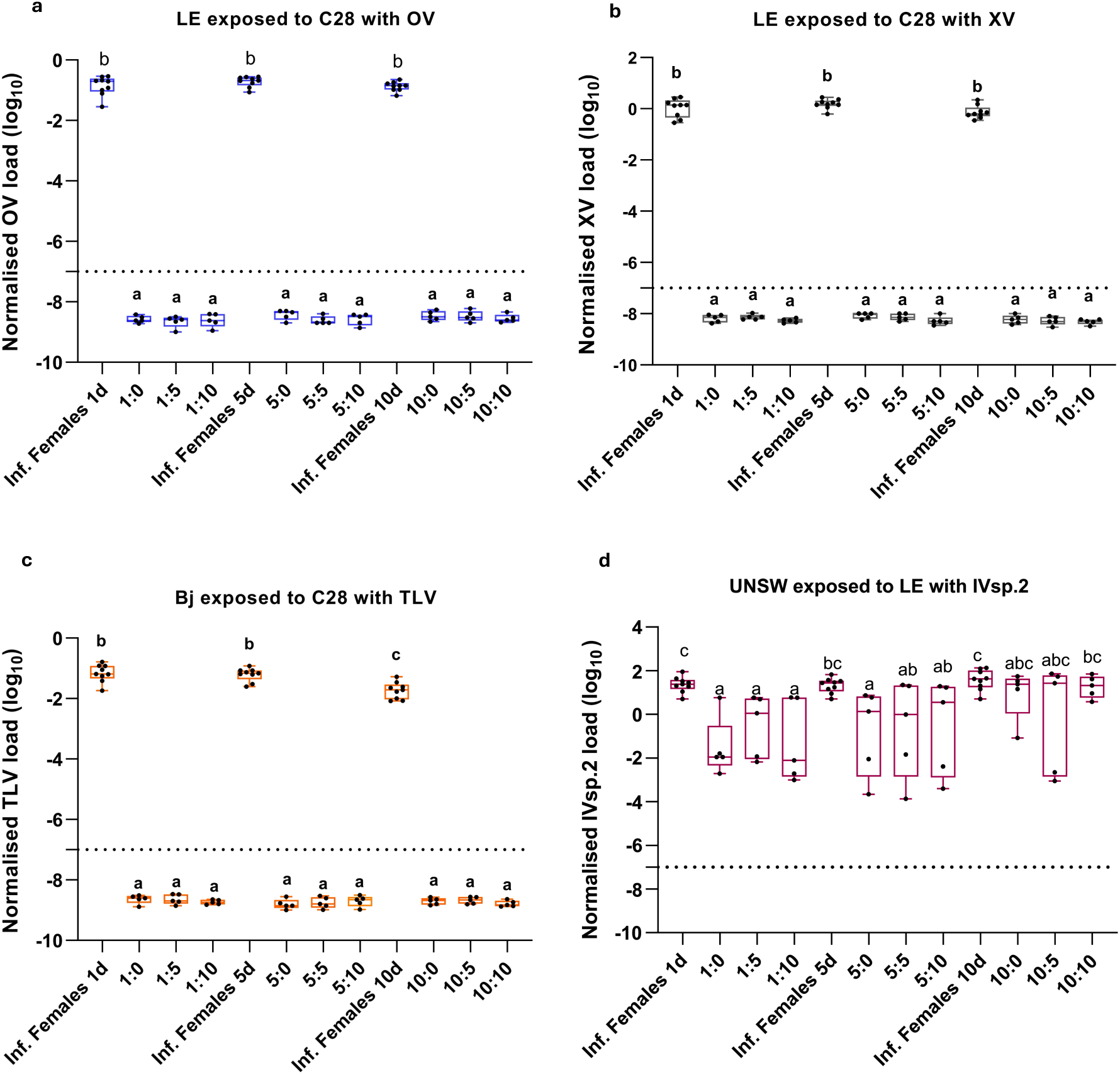
Normalised viral load (log_10_) in females of infected (n=9) and uninfected (n=5 for each time point) laboratory populations after different cohabitation and incubation times. (a) OV and (b) XV load after cohabitation of infected C28 and uninfected LE individuals followed by incubation; (c) TLV load after cohabitation of infected C28 and uninfected Bj individuals followed by incubation; (d) IVsp.2 load after cohabitation of infected LE and uninfected UNSW individuals followed by incubation. The virus detection threshold was set at 1e-7 virus-specific RNA-dependent polymerase (RdRp) or nucleocapsid (N) per *ef1a* transcript, shown by the horizontal dashed line. Means that differ significantly from each other are represented by different letters for each population separately.

Notably, a different infection pattern was observed for IVsp.2. After 1 day cohabitation, IVsp.2 load in females of infected LE was significantly higher (Kruskal-Wallis chi-squared = 34.58, df = 11, *p* < 0.01) than in exposed females of uninfected UNSW (Figure 3d). However, as the cohabitation time increased, the IVsp.2 load in uninfected females also increased and finally, after 5 days cohabitation and 5 days incubation (5:5) and after that (5:10, 10:0, 10:5, 10:10), no significant difference in IVsp.2 load between females of the infected LE population and exposed females of the uninfected UNSW population was observed (Figure 3d). No significant differences among exposed females were observed across the different incubation times after 1 day (1:0, 1:5 and 1:10), 5 days (5:0, 5:5 and 5:10) and 10 days cohabitation (10:0, 10:5), but after 10 days of incubation following 10 days cohabitation (10:10), the IVsp.2 load was significantly increased in exposed females compared to the other incubation times (1:0, 1:5, 1:10, 5:0).

### 3.4 Viral load in different tissues

All four viruses were detected in both somatic and reproductive tissues of adult flies and in most cases, viral load differed significantly between fly types (yeast-fed females, not yeast-fed females and not yeast-fed males) in both GOS and HAC (*p* < 0.01) (Figures 4a–d; Table S4). In GOS females, OV load appeared to be higher in the ovaries of yeast-fed females compared to the other tissues of yeast-fed females, although a significant difference was observed only between the ovaries and salivary glands. In contrast, in not yeast-fed females no significant differences in OV load were observed among tissues (Figure 4a). In GOS males, OV load was significantly higher in the brain compared to other tissues, and higher in thorax compared to salivary glands and testes. In HAC, however, the OV load was very low and close to the detection limit. Nevertheless, the tissue tropism trend observed in HAC was similar to the patterns of OV load seen in GOS.

**Figure 4.**
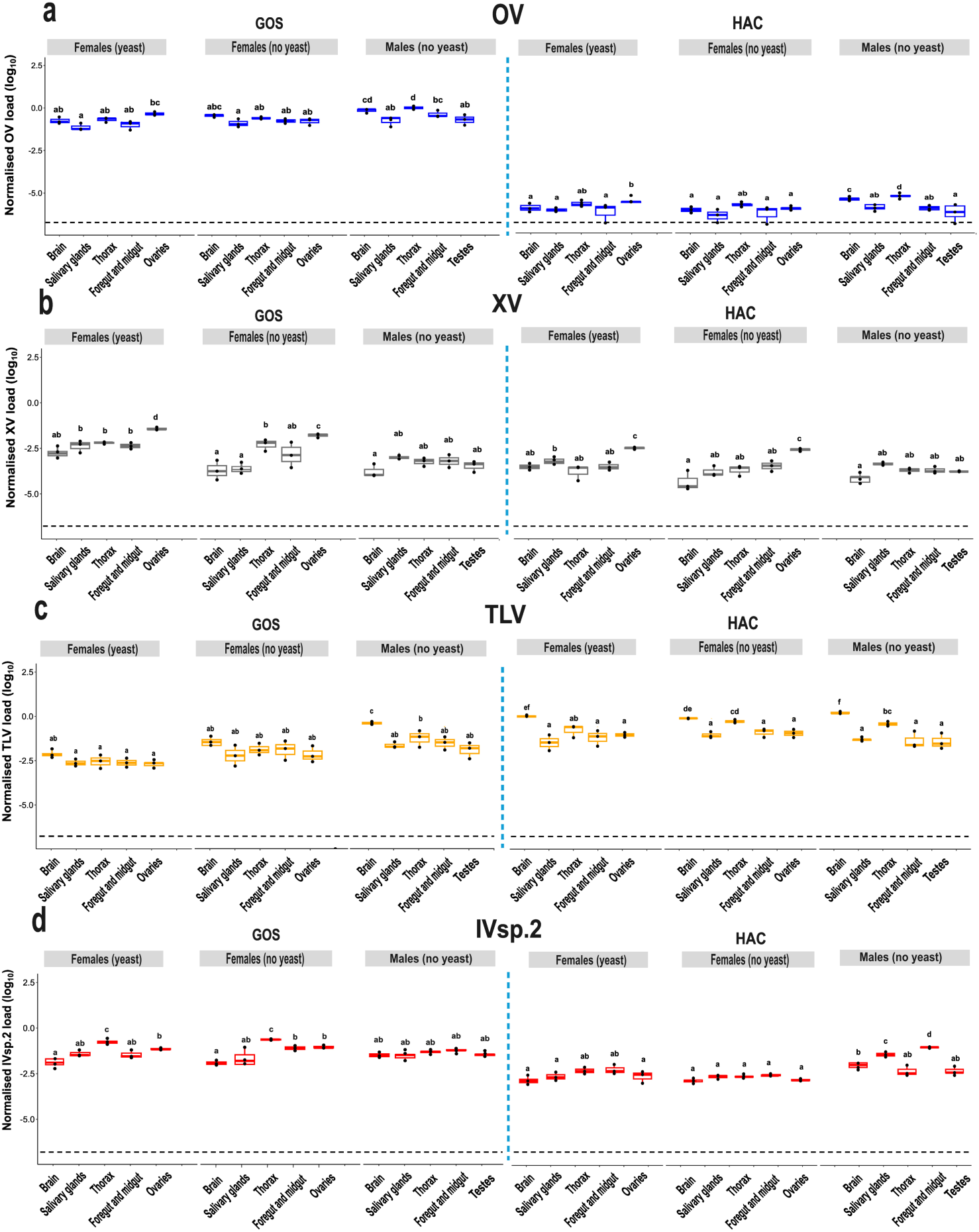
Normalised viral load (log_10_) of (a) Bactrocera tryoni orbivirus (OV), (b) Bactrocera tryoni xinmovirus or *Trocevirus haikouense* (XV), (c) Bactrocera tryoni toti-like virus (TLV) and (d) Bactrocera tryoni iflavirus sp. 2 (IVsp.2) in brain, salivary glands, thorax, foregut and midgut, and ovaries or testes in females (yeast-fed), females (not yeast-fed) and males (not yeast-fed) of GOS and HAC. The virus detection threshold was set at 1e-7 virus-specific RNA-dependent polymerase (RdRp) or nucleocapsid (N) per *ef1a* transcript, shown by the horizontal dashed line. Means that differ significantly from each other are represented by different letters for each population separately. The different tissue types are listed (left to right) in the order of dissection. No consistent pattern indicative of dissection-related leakage of virus across tissues was observed.

In yeast-fed and not yeast-fed GOS and HAC females XV was predominantly found in the ovaries that had significantly higher viral load compared to other tissues (Figure 4b). Additionally, XV load was significantly higher in ovaries of yeast-fed females than not yeast-fed females in GOS, whereas no such difference was observed in HAC. However, there was no significant difference in XV across different tissues of GOS and HAC males (Figure 4b).

In HAC, TLV showed significantly higher loads in the brain and thorax of not yeast-fed females and in the brain of yeast-fed females compared to other tissues in females. In contrast, TLV did not differ significantly across different tissues of yeast-fed and not yeast-fed females in GOS. Additionally, TLV was significantly higher in the brain and thorax of HAC males and in the brain of GOS males compared to other tissues of males (Figure 4c).

In GOS, IVsp.2 load was significantly higher in the thorax of females compared to other tissues of females and males, while no significant differences were observed across tissues of GOS males. In contrast, IVsp.2 load in HAC was significantly higher in the thorax, foregut and midgut of males compared to other tissues of females and males, whereas no significant differences were observed across all tissues in HAC females (Figure 4d).

### 3.5 Viral load across developmental stages

All four viruses, OV (Figure 5a), XV (Figure 5b), TLV (Figure 5c) and IVsp.2 (Figure 5d) varied in their load across developmental stages in GOS and HAC (*p* < 0.01) except for TLV in HAC (*p* = 0.695) (Figure 5c; Table S5). A clear distinction was observed among viruses: OV, XV and TLV were detected across all developmental stages, whereas IVsp.2 was barely detectable in eggs, larvae and pupae. For some viruses the viral load was low, e.g. the OV load in HAC was overall low (Figure 5a), and the IVsp.2 load in pre-adult stages was also low (close to the detection threshold) and differed significantly from adult females and males in both GOS and HAC. In contrast, the viral load of OV in GOS, and XV and TLV in both GOS and HAC, were high in pre-adult stages (eggs, larvae and pupae) and adults (males and females). The viral load of the four viruses varied between males and females across the two fly populations, with significant differences observed in OV load in GOS and HAC, XV load in HAC (but not GOS), TLV load in GOS (but not HAC) and IVsp.2 load in both GOS and HAC. In contrast, no significant differences were found in TLV and IVsp.2 load for pre-adult stages in neither GOS nor HAC. Similarly, the viral load remained unchanged during the larval and pupal stages for XV in GOS and HAC, and for OV in HAC but not GOS.

**Figure 5.**
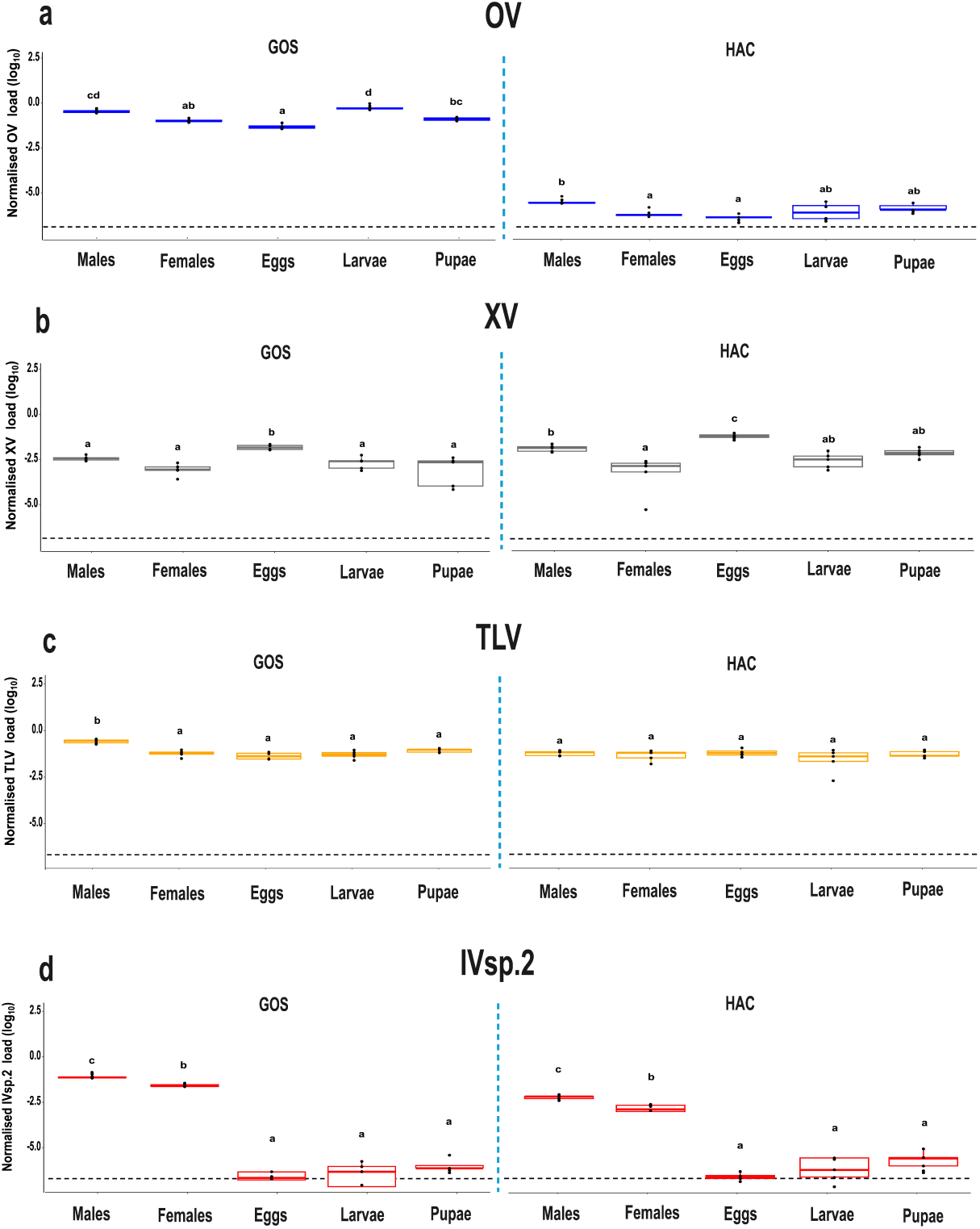
Normalised viral load (log_10_) of (a) Bactrocera tryoni orbivirus (OV), (b) Bactrocera tryoni xinmovirus or *Trocevirus haikouense* (XV), (c) Bactrocera tryoni toti-like virus (TLV) and (d) Bactrocera tryoni iflavirus sp. 2 (IVsp.2) in males, females, eggs, larvae and pupae of GOS and HAC. The virus detection threshold was set at 1e-7 virus-specific RNA-dependent polymerase (RdRp) or nucleocapsid (N) per *ef1a* transcript, shown by the horizontal dashed line. Means that differ significantly from each other are represented by different letters for each population separately.

## 4. Discussion

Our comparative study of four RNA viruses in *B. tryoni* revealed that OV, XV, TLV and IVsp.2 exhibit distinct transmission routes, tissue tropism and infection patterns across developmental stages. Vertical transmission of OV, XV and TLV is supported by (i) their consistent detection within and/or on eggs, (ii) their presence across all developmental stages, and (iii) the absence of horizontal transmission in cohabitation experiments. In contrast, IVsp.2 is horizontally transmitted, as evidenced by (i) its predominant absence from eggs, (ii) its very low viral load in other pre-adult stages, and (iii) its consistent horizontal transmission in cohabitation experiments. Tissue profiling of all four viruses revealed infection in both somatic and reproductive tissues, with significant enrichment of XV in ovaries and TLV in the brain and thorax, particularly in HAC, whereas IVsp.2 generally showed low viral load in the brain and higher loads in the digestive tissues, particularly in HAC males. This supports the finding of the horizontal transmission mode of IVsp.2, as well as the potential for its host effects that may be distinctly different from those of the three vertically transmitted viruses.

Together with previous findings about the transmission modes of CV, IVsp.1 and SV in *B. tryoni* (Morrow et al., 2023; Pradhan et al., 2026) our study has established a comprehensive transmission mode analysis of seven viruses of *B. tryoni* which include almost all known viruses of the virome of this host species (Table 1). Such detail exists for few insect species such as honey bee, *D. melanogaster* and mosquitoes (Chen et al., 2006; Webster et al., 2015; Webster et al., 2016; Bonning, 2019), as well as other tephritid pest species such as *C. capitata* (Sharpe et al., 2021; Hernández-Pelegrín et al., 2024a; Hernández-Pelegrín et al., 2024b). Our and previous studies of the viruses in *B. tryoni* (Morrow et al., 2023; Sharpe et al., 2024; Pradhan et al., 2026), reveal that the majority of investigated RNA viruses (5 of 7) in *B. tryoni* are vertically transmitted (Table 1). This challenges the perhaps prevailing view that viruses in a host are mostly horizontally transmitted (Fermin, 2018), as seen for example for plant viruses in plants (García-Ordóñez and Pagán, 2024), with important consequences for the understanding of the evolution of host–virus interactions in insects. It has been hypothesized that horizontal and vertical transmission result in different selective pressures on the virulence and the dynamics of host–pathogen interactions (Ewald, 1987; Lipsitch et al., 1996; García-Ordóñez and Pagán, 2024). Horizontally transmitted viruses are expected to be more virulent and to occur at higher viral loads, as greater pathogen output increases horizontal transmission success. In contrast, vertically transmitted viruses rely on host survival and reproduction for transmission, so they may have lower virulence and persist as covert infections, as any reduction in host survival or reproduction would limit their persistence; they may even have beneficial effects on the host (Ewald, 1987; Lipsitch et al., 1996; García-Ordóñez and Pagán, 2024).

**Table 1.**
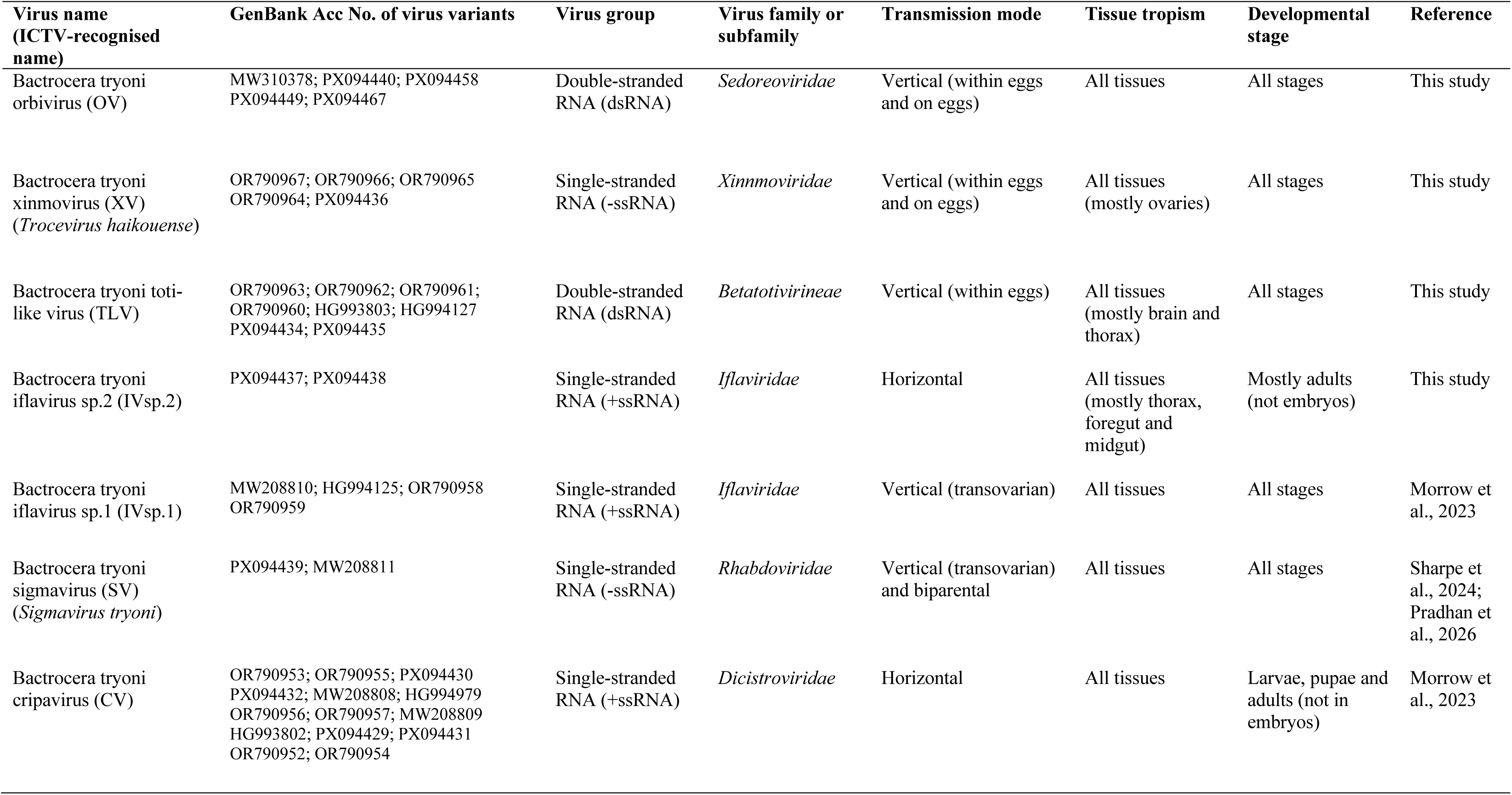
List of seven RNA viruses and their transmission modes, tissue tropism and localisation across developmental stages in *Bactrocera tryoni*. Our study investigated the transmission mode and viral load across tissues and developmental stages for four of these viruses. The parameters for the three other viruses are listed with the references to previous studies. For two of the seven viruses, the International Committee on Taxonomy of Viruses (ICTV) has provided recognised virus names.

### 4.1 Vertical transmission

The egg testing and surface bleaching assays revealed transmission within and on eggs for OV and XV, and within eggs for TLV. Related viruses of these virus families are also vertically transmitted in their insect hosts. For instance, members of the family *Xinmoviridae*, such as *Aedes anphevirus* (AeAV), are maintained through vertical transmission in mosquito populations (Parry and Asgari, 2018). Likewise, toti-like viruses are vertically transmitted in *Aedes aegypti*, with detection in ovaries and maintenance across multiple generations (Coatsworth et al., 2022). Furthermore, the detection of OV, XV and TLV in ovaries and testes suggests maternal and/or paternal transmission are both possible for these viruses. The type of vertical transmission of OV, XV and TLV should be investigated further by testing eggs obtained from infected females mated with uninfected males and eggs of the reciprocal mating combination, to ascertain whether the vertical transmission is maternal and/or paternal, respectively.

### 4.2 Horizontal transmission

The cohabitation experiment revealed the absence of horizontal transmission for OV, XV and TLV. It is noted, however, that we used *B. tryoni* and *B. jarvisi* to investigate the potential for horizontal transmission of TLV, since all *B. tryoni* laboratory populations were infected with TLV. Therefore, absence of TLV horizontal transmission could also indicate narrow host specificity; yet *B. jarvisi* is a host for a genetically similar virus (98.23% nucleotide similarity of the RdRp gene) (Sharpe et al., 2021), and *B. tryoni* and *B. jarvisi* are closely enough related to still hybridise under laboratory conditions (Morrow et al., 2014).

In contrast, IVsp.2 exhibited horizontal transmission. Following cohabitation with infected LE females, IVsp.2 was detected in exposed females of the uninfected UNSW population, most strongly after 10 days of cohabitation. With increasing cohabitation duration, IVsp.2 was detected in exposed females, suggesting that prolonged contact with infected females, shared food and water, or exposure to excretions of infected individuals, facilitated horizontal transmission of IVsp.2 in *B. tryoni*, similar to what has been reported for CV in *B. tryoni* (Morrow et al., 2023), nodavirus and noravirus in *C. capitata* (Hernández-Pelegrín et al., 2024b), and Drosophila C virus and noravirus in *Drosophila* (Gupta et al., 2017; Habayeb et al., 2009).

IVsp.2 load increased across cohabitation periods and the IVsp.2 load of exposed and infected flies was indistinguishable after the 10 day cohabitation period. This pattern implies that while infection may occur early, viral replication progresses over time resulting in a statistically significant increase in IVsp.2 load. Therefore, while OV, XV and TLV showed no detectable horizontal transmission under our experimental conditions, IVsp.2 clearly exhibited this capacity. This finding aligns with broader patterns of horizontal transmission generally observed for viruses of the genus *Iflavirus*.

Transmission patterns within the *Iflaviridae* vary among virus species, with examples of both horizontal and vertical transmission across the family. Several iflaviruses adopt a mixed transmission mode strategy; however, the efficiency of the horizontal versus vertical transmission pathways is not always equal, with some viruses showing a clear bias towards one transmission mode over the other (Mattia et al., 2025). For instance, IVsp.1 is predominantly transmitted vertically, from adults to offspring in *B. tryoni* (Morrow et al., 2023). Similarly, *C. capitata* has two vertically transmitted iflaviruses (CcaIV2 and CcaIV4), and, based on egg sterilisation and mating experiments, CcaIV2 and CcaIV4 were found to be primarily transmitted vertically (maternally). Moreover, CcaIV4 showed evidence of limited horizontal transmission (Hernández-Pelegrín et al., 2024b). An iflavirus was also detected in *Glossina morsitans morsitans* and its tissue tropism suggests both horizontal and vertical transmission (Meki et al., 2021). In *Spodoptera exigua*, the two different iflaviruses SeIV-1 and SeIV-2 display contrasting transmission strategies. After inoculation of *S. exigua* egg masses with SeIV-1 and SeIV-2, SeIV-1 loads increased and remained high across larval, pupal and adult stages, supporting horizontal transmission via faeces or regurgitation. In contrast, SeIV-2 maintained low viral loads, suggesting a persistent infection that is likely maintained through vertical transmission (Carballo et al., 2020). This is consistent with our results showing different transmission modes for the two phylogenetically diverged iflaviruses IVsp.1 and IVsp.2 in *B. tryoni.* In comparison, sacbrood virus (SBV) of honey bee has also both vertical and horizontal transmission, as it occurs among adult bees and from adult workers to larvae through contaminated food resources such as brood provisioning: honey, pollen and royal jelly (Shen et al., 2005). Furthermore, Helicoverpa armigera iflavirus (HaIV) was found to use both horizontal and vertical transmission routes; its peroral horizontal transmission was highly efficient and dose-dependent, resulting in 100% infection, whereas vertical transmission occurred at a much lower rate of <28.57% (Yuan et al., 2017).

### 4.3 Virus localisation across tissues

The four viruses exhibited distinct patterns of tissue localisation. The vertically transmitted viruses OV, XV and TLV were distributed across reproductive and somatic tissues, indicating systemic infection and potential adaptation for reliable vertical transmission across generations. Although IVsp.2 was detected in different tissues, it appeared more strongly associated with digestive tissues, particularly in males of HAC, supporting the horizontal transmission of this virus in the cohabitation experiment. OV abundance appeared to increase in the ovaries of yeast-fed females, although this increase was not statistically significant. OV load may be associated with the development of reproductive tissues, a process that depends on yeast (protein) consumption by female fruit flies. By comparison, XV was detected in ovaries regardless of yeast consumption, indicating that its replication in reproductive tissues is not dependent on yeast-triggered ovarian development. TLV load was high in the brain and thorax, particularly in HAC, leading to the possibility that, if an interaction with the nervous system and flight muscles occurs, this could influence key behaviours such as mating and dispersal. Together, these results show that vertically transmitted viruses in *B. tryoni* are not confined to reproductive or digestive tissues. Similar broad tissue tropism has been documented for vertically transmitted iflaviruses: Antheraea mylitta iflavirus, which was present in the fat body, midgut, Malpighian tubules, silk glands and trachea of the tropical tasar silkworm *Antheraea mylitta* (Ponnuvel et al., 2022), Lymantria dispar iflavirus 1 which was detected in the fat body, midgut, ovaries and haemocytes of the spongy moth *Lymantria dispar* (Carrillo-Tripp et al., 2014), and CcaIV2 and CcaIV4 which were present in the gut, crop, legs, brain, ovaries and testes of *C. capitata* strain Vienna 8A (Hernández-Pelegrín et al., 2024b) suggesting that widespread infection of host tissues may be a common feature of vertically transmitted iflaviruses. In contrast to the three vertically transmitted viruses of *B. tryoni* analysed in our study, the high viral load of IVsp.2 in digestive tissues, particularly in HAC males, suggests that this virus relies on horizontal transmission for its spread. This pattern aligns with reports of high viral loads of horizontally transmitted nodavirus and noravirus in digestive tissues such as the gut and crop of *C. capitata* (Hernández-Pelegrín et al., 2024b). Similarly, larval infection via the oral route has been demonstrated when purified *C. capitata* noravirus was incorporated into larval diet (Hernández-Pelegrín et al., 2024a), supporting the role of the digestive tract as a key entry point for horizontally transmitted viruses. However, our study also demonstrated that the horizontally transmitted IVsp.2 was not restricted to the digestive tract but was also detected in other tissues, such as ovaries and testes, albeit at lower loads. It is possible that IVsp.2 may not be integrated into developing oocytes and only be present in ovarian tissues. Similarly, it may not be transmitted paternally via sperm. Therefore, further targeted studies are required to investigate this.

### 4.4 Virus load across developmental stages

When looking at developmental stages, OV, XV and TLV were detected in all stages, yet the patterns diverged. For OV, XV and TLV, presence in eggs reaffirmed that these viruses are vertically transmitted. TLV load remained relatively stable across all stages, while OV and XV showed fluctuations perhaps reflecting differences in replication rates or host immune responses against these viruses at particular developmental stages. IVsp.2, however, exhibited a clear difference: it was almost undetectable in eggs and pre-adult stages but had a significantly higher viral load in adults. This confirms the cohabitation results, where IVsp.2 load showed a significant increase in exposed flies after a 10 day cohabitation and incubation period (10:10), with a consistent upward trend of increasing load over time, indicating that an extended cohabitation period after initial exposure is required for reaching a high viral load. These patterns are consistent with previous findings in *C. capitata*, where the vertically transmitted CcaIV2 and CcaIV4 were detected across the larval, pupal and adult stages of the Vienna 8A fly strain, whereas the horizontally transmitted nodavirus was mainly found in adults of the Madrid fly strain (Hernández-Pelegrín et al., 2024b). Similar stage-specific infection dynamics have been reported for Solenopsis invicta virus 3 (*Solinviviridae*) where viral genome accumulation increases over time in adults but remains low in larval stages (Valles et al., 2014).

### 4.5 Unknown virus-virus interactions and host effects

Our study was conducted on fly populations carrying multiple viruses and, thereby, results may also be influenced by virus-virus interactions. Coinfections can potentially have additive, synergistic and antagonistic effects on transmission modes and efficiencies, tissue tropism and developmental stage distribution, and this requires further investigation. Based on recent studies, no coinfection between the vertically transmitted IVsp.1 and SV was observed in *B. tryoni* field and laboratory populations (Sharpe et al., 2024). This absence may be due to their low overall prevalence in field populations, their mostly maternal transmission (Morrow et al., 2023), less effective paternal than maternal transmission of SV (Pradhan et al., 2026), potential competitive exclusion and/or immune priming effects (Sharpe et al., 2024). The investigation of virus-virus interactions are important, as some studies have observed that insect-associated viruses in mosquitoes can negatively (Vasilakis and Tesh, 2015; Patterson et al., 2020; Altinli et al., 2021) or positively (Olmo et al., 2023) affect the replication and transmission efficacy of several arboviruses.

RNA viruses related to OV, XV, TLV and IVsp.2 have been identified in tephritid fruit flies and other Diptera, and across a wide range of insect orders (Shi et al., 2016; Qi et al., 2023). *Sedoreoviridae* (Reovirales) similar to OV have been reported in tephritids, such as *B. dorsalis* and *Zeugodacus cucurbitae* (Zhang et al., 2022; Pradhan et al., 2024a), as well as in biting midges and mosquitoes (Colmant et al., 2017), and in bat flies (Ramírez-Martínez et al., 2021). A virus related to XV, *Trocevirus haikouense* (>89% amino acid similarity with XV) has been found in *B. dorsalis* (Sharpe et al., 2021), and several viruses of the *Xinmoviridae* (Mononegavirales) have been reported from the bugs *Cletus punctiger* and *Zicrona caerulea* (Divekar et al., 2024) and multiple insect orders (Hernández-Pelegrín, 2025). Various *Betatotivirineae* (Ghabrivirales) similar to TLV have been detected in tephritids, including *B. jarvisi*, *B. dorsalis* and *Z. cucurbitae* (Sharpe et al., 2021; Pradhan et al., 2024a), as well as in the rice planthoppers *Nilaparvata lugens*, *Laodelphax striatellus* and *Sogatella furcifera* (Huang et al., 2023). Various *Iflaviridae* (Picornavirales) have been reported in *C. capitata* (Llopis-Gimenez et al 2017, Hernández-Pelegrín et al., 2022) and other insects including the splayed deer fly *Chrysops caecutiens*, the horse fly *Hybomitra stigmoptera* and the thrips *Haplothrips nigricornis* (Litov et al., 2023). However, sequence similarities are low compared with the IVsp.2 investigated in this study.

The effects of many insect-associated viruses detected in transcriptomes on host biology and fitness remain poorly understood. This knowledge gap remains because RNA viruses often cause covert infections (particularly in laboratory populations), lacking visible symptoms. Sometimes it is also difficult to evaluate the effects of viruses because they are omnipresent in laboratory populations (Morrow et al., 2023). Nevertheless, growing evidence suggests that some RNA viruses can impose sublethal fitness costs. For instance, IVsp.1 (in coinfection with CV and other viruses) has been linked to slower development and reduced survival in *B. tryoni* under stress (Sharpe et al., 2026). SV causes paralysis in *Drosophila* and *B. tryoni* when exposed to CO₂ at low temperatures (Longdon et al., 2012; Pradhan et al., 2026). These findings underscore the need for further research to uncover the hidden biological consequences of RNA virus infections in insects.

## 6. Conclusions

Our study provides critical insights into the transmission modes and tissue and developmental stage-specific localisation of OV, XV, TLV and IVsp.2 in *B. tryoni*. By combining egg testing and bleaching, and cohabitation experiments with detailed analyses of tissue- and stage-specific viral load, we demonstrated that OV, XV and TLV are vertically but not horizontally transmitted, while IVsp.2 relies on horizontal transmission and replicates primarily in adult flies. Our study, in combination with previous research, shows that the majority of the known (including the most common) RNA viruses in *B. tryoni* have vertical transmission.

Persistent infections caused by vertically or horizontally transmitted viruses may reduce the efficiency of mass production and compromise the quality of mass-reared individuals for pest management approaches such as SIT. Our study addresses an important knowledge gap in understanding how these viruses spread and persist in both field and laboratory populations and may also inform the assessment of their potential impacts on host biology and fitness. Furthermore, the findings of our study may contribute to the development of targeted biological control strategies that involve viruses (Pradhan et al., 2024b; Coffman, 2025).

## Supporting information

Excel file with viral load data

## Ethics approval

*Bactrocera tryoni* is a common pest insect and research on it does not require animal ethics approval. Irrespective of this, flies were anaesthetised before dissection and nuclear acid extraction.

## Author contributions

The study was conceived and designed by FB, MR, and JLM. Data collection was performed by FB and analyses were conducted by FB with the help of SKP and under the supervision of MR and JLM. FB wrote the manuscript with feedback from MR and JLM. All authors agreed to the submission of the manuscript.

## Acknowledgements

We thank Geraldine Tilden and Alihan Katlav for their help with fly maintenance and statistical analyses.

## Data availability statement

All data are contained within the article, the Supplementary Tables S1-S5 and the Supplementary File (which also contains all qPCR data).

