## Supplementary Tables S1-5 for "Comparative studies of transmission mode and localisation patterns of common RNA viruses in Queensland fruit fly (*Bactrocera tryoni*) reveal most are vertically transmitted"

**Table S1.** Laboratory populations of *Bactrocera tryoni* and *Bactrocera jarvisi* used in this study

| **Fly population** | **Establishment date** | **Reference** |
|---|---|---|
| *B. tryoni* GOS | 2009 | Langford et al. 2014 |
| *B. tryoni* HAC | 2009 | Morrow et al. 2015 |
| *B. tryoni* lemon eyes LE | 2001 | Zhao et al. 2003 |
| *B. tryoni* UNSW | 2009 | Sharpe et al. 2021 |
| *B. tryoni* C28 | 2017 | Morrow et al. 2023 |
| *B. jarvisi* Bj | 2010 | Morrow et al. 2014; Morrow et al. 2015 |

**Table S2.** Primers used in this study

| Target | Primer name | Primer sequence |
|---|---|---|
| Orbivirus | BtOVS1 primer set 1 | Forward: CGATTATGCTTTCGGCGTGG |
| | | Reverse: TCTCACCTGACCCTTCGCTA |
| Xinmovirus | BtXV primer set 2 | Forward: ACCTTTCTACTAACCGAGGCAC |
| | | Reverse: TGCAGTCTGGACACACGATT |
| Toti-like virus | BtTLV primer set 1 | Forward: CGGCAGGGTATGCACTATGT |
| | | Reverse: GGTAACCCTTCTAAGCGCCA |
| Iflavirus sp. 2 | BtIVsp.2 primer set 2 | Forward: TCGCAAGGAGGAACACGTTTA |
| | | Reverse: AGAAGCCATAAGCTCTCGCAA |
| Iflavirus sp. 1 | BtIVsp.1 primer set 1 | Forward: TCCAGTCTTGGACTTTGAGGA |
| | | Reverse: GGTGCTAAATTCCATTGCATC |
| Sigmavirus | BtSVN primer set 1 | Forward: CCTACATCCAGAGAAACC |
| | | Reverse: GTGAACCTACAACTTTTGTC |
| Cripavirus | BtDV1 primer set 1 | Forward: CCCACAACGCAACATCATGT |
| | | Reverse: TGTGGACCCGCGTAATAACA |
| *Bactrocera tryoni* | Btef1a | Forward: TGTTCAAGTCTGTTGGCGGAT |
| | | Reverse: GTGGTGGTCGACTTTCCAGA |

**Table S3.** Statistical data on the prevalence of seven RNA viruses in various laboratory populations

| **Population name** | **Statistic parameters** |
|---|---|
| GOS | Cochran's Q: 88.035, df: 6, n=20, $p < 0.01$ |
| HAC | Cochran's Q: 80.947, df: 6, n=20, $p < 0.01$ |
| LE | Cochran's Q: 94, df: 6, n=20, $p < 0.01$ |
| Bj | Cochran's Q: 104.818, df: 6, n=20, $p < 0.01$ |
| C28 | Cochran's Q: 99.736, df: 6, n=20, $p < 0.01$ |
| UNSW | Cochran's Q: 55.831, df: 6, n=20, $p < 0.01$ |

**Table S4.** Statistical data on the tissue-specific viral loads of four RNA viruses

| **Virus and Population name** | **Statistic parameters** |
|---|---|
| OV (GOS) | $F_{14, 45} = 37.735$, $p < 0.01$ |
| OV (HAC) | $F_{14, 45} = 37.627$, $p < 0.01$ |
| XV (GOS) | $F_{14, 45} = 40.035$, $p < 0.01$ |
| XV (HAC) | $F_{14, 45} = 35.49$, $p < 0.01$ |
| TLV (GOS) | $F_{14, 45} = 38.566$, $p < 0.01$ |
| TLV (HAC) | $F_{14, 45} = 35.567$, $p < 0.01$ |
| IVsp.2 (GOS) | $F_{14, 45} = 40.831$, $p < 0.01$ |
| IVsp.2 (HAC) | $F_{14, 45} = 40.61$, $p < 0.001$ |

**Table S5.** Statistical data of four RNA viruses load in different developmental stages

| Virus and Population name | Statistic parameters |
|---|---|
| OV (GOS) | $F_{4,24} = 18.288$, $p < 0.01$ |
| OV (HAC) | $F_{4,24} = 14.33$, $p < 0.01$ |
| XV (GOS) | $F_{4,24} = 16.649$, $p < 0.01$ |
| XV (HAC) | $F_{4,24} = 19.841$, $p < 0.01$ |
| TLV (GOS) | $F_{4,24} = 15.396$, $p < 0.01$ |
| TLV (HAC) | $F_{4,24} = 2.217$, $p = 0.695$ |
| IVsp.2 (GOS) | $F_{4,24} = 19.755$, $p < 0.01$ |
| IVsp.2 (HAC) | $F_{4,24} = 16.352$, $p < 0.01$ |
